# The gliopeptide ODN, a ligand for the benzodiazepine site of GABA_A_ receptors, boosts functional recovery after stroke

**DOI:** 10.1101/2020.03.05.977934

**Authors:** Rhita Lamtahri, Mahmoud Hazime, Emma K Gowing, Raghavendra Y. Nagaraja, Julie Maucotel, Michael Alasoadura, Pascale Quilichini, Katia Lehongre, Benjamin Lefranc, Katarzyna Gach-Janczak, Ann-Britt Marcher, Susanne Mandrup, David Vaudry, Andrew N. Clarkson, Jérôme Leprince, Julien Chuquet

## Abstract

Following stroke, the survival of neurons and their ability to re-establish connections is critical to functional recovery. This is strongly influenced by the balance between neuronal excitation and inhibition. In the acute phase of experimental stroke, lethal hyperexcitability can be attenuated by positive allosteric modulation of GABA_A_ receptors (GABA_A_R). Conversely, in the late phase, negative allosteric modulation of GABA_A_R can correct the sub-optimal excitability and improves both sensory and motor recovery. Here, we hypothesized that octadecaneuropeptide (ODN), an endogenous allosteric modulator of the GABA_A_R synthesized by astrocytes, influences the outcome of ischemic brain tissue and subsequent functional recovery. We show that ODN boosts the excitability of cortical neurons, which make it deleterious in the acute phase of stroke. However, if delivered after day 3, ODN is safe and improves motor recovery over the following month in two different paradigms of experimental stroke in mice. Furthermore, we bring evidence that during the sub-acute period after stroke, the repairing cortex can be treated with ODN by means of a single hydrogel deposit into the stroke cavity.

**SIGNIFICANCE STATEMENT:** Stroke remains a devastating clinical challenge because there is no efficient therapy to either minimize neuronal death with neuroprotective drugs or to enhance spontaneous recovery with neurorepair drugs. Around the brain damage, the peri-infarct cortex can be viewed as a reservoir of plasticity. However, the potential of wiring new circuits in these areas is restrained by a chronic excess of GABAergic inhibition. Here we show that an astrocyte-derived peptide (ODN), can be used as a delayed treatment, to safely correct cortical excitability and facilitate sensorimotor recovery after stroke.

## 1 Introduction

Stroke remains a devastating clinical challenge because there is no efficient therapy to either minimize neuronal death with neuroprotective drugs or to enhance spontaneous recovery with neurorepair drugs. The chronic and ever-changing balance between neuronal excitation and inhibition is a primary trigger for the initial stroke progression as well as the impairment in the ability to regain function. Given its predominant role in the control of inhibition, GABAergic transmission remains a major target to act on the dynamic of this balance (Bachtiar and Stagg, 2014; Roux and Buzsáki, 2015). The use of pharmacological manipulations targeting GABA_A_ receptors (GABA_A_R) to modify the course of ischemic cell damage and their sequelae is an old quest: numerous pre-clinical experiments have demonstrated that sustaining GABA neurotransmission, to counteract the initial excitotoxic effect of glutamate released minutes to hours after the onset of stroke can afford protection (Sydserff *et al*., 1995; Shuaib and Kanthan, 1997; Green *et al*., 2000; Schwartz-Bloom and Sah, 2001; Marshall *et al*., 2003). In this respect, benzodiazepines with positive allosteric modulation (PAM) profile were repeatedly shown to be neuroprotective after cerebral ischemia (Schwartz *et al*., 1994,1995; Schwartz-Bloom *et al*., 1998; Galeffi *et al*., 2000). However, like all neuroprotective drug therapies to-date, positive GABA modulators have failed to translate into clinical use.

In recent years, GABA modulation in stroke has been revitalized with the understanding that once the infarction is consolidated, the excitation and inhibition balance switches from hyperexcitability to hypoexcitability in the peri-infarct cortex, due to a loss of the astroglial GABA tranporter, GAT3, and resultant excess in ambient GABA (Clarkson *et al*., 2010; Carmichael, 2012). This prolonged synaptic depression is thought to limit neuronal circuit reorganization and therefore the regain of sensorimotor functions (Hummel *et al*., 2009; Kim *et al*., 2014). Dampening the stroke-induced elevation in inhibition using a negative allosteric modulator (NAM) targeting the benzodiazepine site of GABA_A_R, enhances sensorimotor recovery in mice and rats (Clarkson *et al*., 2010, 2015; Lake *et al*., 2015; Alia *et al*., 2016; Orfila *et al*., 2019).

Endozepines, known as the endogenous ligands of benzodiazepine-binding sites, comprise the diazepam binding inhibitor (DBI, also known as acyl-CoA binding protein, ACBP) and its processing products, including the octadecaneuropeptide (ODN), a small peptide of 18 amino-acids (DBI_33-50_) (Tonon *et al*., 2020). In the brain, DBI is one of the major proteins expressed and released by astrocytes but not by neurons (Loomis *et al*., 2010; Tonon *et al*., 2020). DBI and ODN bind to the benzodiazepine site of the GABA_A_R where they act as allosteric neuromodulators (Bormann, 1991; Barmack *et al*., 2004; Quian *et al*., 2008; Möhler, 2014; Dumitru *et al*., 2017). Point mutation targeting the benzodiazepine site of GABA_A_R renders neuronal cells insensitive to ODN (Dumitru *et al.,* 2017). At micromolar concentrations, electrophysiological studies show that ODN acts as a NAM on GABA_A_Rs, i.e. reducing GABA_A_R mediated inhibition (Guidotti *et al*., 1983; Ferrero *et al*., 1986; Barmack *et al*., 2004; Alfonso *et al*., 2012; Dumitru *et al*., 2017) with no epileptogenic effect (Vezzani *et al*., 1991). ODN appears to be a relatively new astroglial modulator of GABA_A_R signaling and should therefore be considered for its potential to correct the imbalance between excitation and inhibition that arises as a consequence of a stroke. Here, we tested the gliopeptide ODN for its potency to enhance recovery after stroke. We show that ODN boosts the excitability of cortical neurons, which make it deleterious in the acute phase of stroke. However, if delivered after day 3, ODN is safe and improves motor recovery in two different models of focal brain ischemia. Furthermore, we bring novel evidence that during the sub-acute period after stroke, the repairing cortex can be treated with ODN by the means of a single hydrogel injection into the stroke cavity allowing for direct targeting of the peri-infarct cortex.

## Materials and methods

### Animals and approvals

Eight to twelve-week-old (20-25 g) male C57BL/6J mice were purchased from Janvier Laboratories. ACBP knockout (ACBP^−/−^ hereinafter referred to as DBI^−/−^) mice were obtained from Prof. Mandrup Laboratory (University of Southern Denmark). Mice were backcrossed to the C57BL/6J Bom Tac strain for ten generations to obtain a congenic background as previously described (Neess et al., 2011). Wild type (DBI^+/+^) and KO (DBI^−/−^) were littermate generated from heterozygote mice. The homozygote DBI^−/−^ mice used in this study were identified by genotyping PCR. Aged female (20±2 months) mice were obtained from the Hercus Taieri Resource Unit at the University of Otago. Experiments, approved by the Ethics Committee for Animal Research of Normandy or the University of Otago Animal Ethics Committee, were conducted by authorized investigators in accordance with the recommendations of the European Communities 86/609/EEC. All procedures were undertaken and reporting done in accordance to the ARRIVE (Animal Research: Reporting *In Vivo* Experiments) guidelines. All *in vivo* procedures were carried out between 8am-6pm in specific experimental rooms.

### *In vivo* electrophysiology

Under isoflurane anesthesia (2-2.5%) two holes were drilled over the whisker barrel cortex with the dura mater intact and two glass micropipettes were inserted for recording and micro-injection. After the surgical procedure, isoflurane was reduced to 1.1±0.1% for a 30-45 min resting period. All *in vivo* recordings were done with an amplifier PowerLab 8/35 and aquired with Labchart software (AD-Instrument).

#### Local field potential (LFP) and extracellular unit recording (EU)

An aCSF-filled glass-micropipette/AgCl/Ag electrode, 3-6 μm diameter opening, was positioned in cortical layer 4 of the whisker barrel cortex. Reference and ground electrodes (AgCl/Ag wire) were inserted into the cerebellum. For ODN microinjection, a glass micropipette (10 μm diameter opening) was placed 50-100 μm away from the recording pipette’s tip. For KCl-induced spreading depolarization waves, 2 μL of KCl (0.5 mol/L) was slowly infused (10 minutes) at a distant site from the recording site (4 mm). The signal was bandpass filtered at 200-2000 Hz and digitized at 20 kHz for EU or at 1-100 Hz and digitized at 4 kHz for LFP. Signals were recorded for 12 min before the microinjection of 1 μg of ODN (0.5 μL) and compared to a 12 min period beginning 3 min after the microinjection ended (3 min). Spike detection and sorting were then performed semi-automatically, using Klusta software suite (Rossant *et al*., 2016), freely available (http://klusta-team.github.io). Spikes were identified from the high frequency component by SpikeDetekt using a thresholding of 4.5 times the standard deviation of the signal. Clustering was performed using Klustakwik to verify the coherence of the clusters of the two periods (pre- vs. post-injection period) and eliminate spikes that are an apparent noise (<1% of total spikes). Waveforms were viewed and extracted using Phy Graphical unit interface. For each animal, the spiking change (%) represent a normalized difference of spikes between the pre- and the post-injection period.

#### Somatosensory evoked potentials (SEP)

The left whiskers were stimulated with a short rod controlled by an Arduino UNO board at 0.1 Hz, 100 times. Signal was acquired at 4 kHz and analysis was performed by a custom-made script in MATLAB, bandpass filtered (1 to 25 Hz) and segmented around each stimulation (−2 to +4 s). Each trial was normalized by subtracting the baseline, i.e signal average of the signal comprised between −1 and −0.1 s. For the SEP slope, 100 trials were superimposed and averaged, and the initial deflection of the LFP was defined as the interval within 20–80% of the peak-to-peak amplitude. The slope was computed by linear regression of the selected region.

### Intravital calcium imaging

A cranial window was made over the somato-sensory cortex as previously described (Chuquet *et al*, 2010). Imaging was performed under anesthesia using 2-photon laser scanning microscope (SP8, Leica). SR-101 (1 μM applied on the cortex for 10 min) and OGB1-AM (1.2μg in 1 μL micro-injected in the cortex) were both excited at 805 nm. Frames (256×256 pixels) were collected at 3 Hz. X-y drift were automatically corrected. All traces were median filtered. Signal was expressed as relative OGB1-AM fluorescence changes (dF/F_0_) where F_0_ is the mean of the lowest 20% of the somatic fluorescence signals. Astrocytic calcium surges were defined as transient increase of dF/F_0_ signal exceeding 3 SDs.

### Stroke models

#### Middle cerebral artery occlusion

Temporary focal cerebral ischemia was induced under general anesthesia (isoflurane 1.5-2%) by occlusion of the right middle cerebral artery (MCAO) using the intraluminal filament technique. Briefly, a nylon thread (80 μm in diameter) with a distal cylinder (1.5 mm and 180 μm in diameter) was inserted into the common carotid artery, advanced to the origin of the MCA (exept for sham) and removed 60 min later to allow reperfusion. A laser-Doppler flowmetry probe (Moor Instruments) was used to continuously monitor CBF. Post-surgery analgesic care comprised intradermal ropivacaine (25 μL at 2.5 mg/mL, Naropeine) around sutures and applications of lidocaine/prilocaine cream (2,5% EMLA, AstraZeneca). The animals were then allowed to recover and were killed at day 2 or 28 after MCAO.

#### Cortical photothrombosis

Focal stroke was induced in the left hemisphere using the photothrombosis method in aged (20±2-month-old) female C57BL/6J mice weighing 35.8±5.6 g (Clarkson *et al*., 2010, 2011, 2019). Briefly, under isoflurane anesthesia (2-2.5% in O_2_) mice were placed in a stereotactic apparatus and the skull was exposed. A cold light source (KL1500 LCD, Zeiss) attached to a 40x objective giving a 2 mm diameter illumination was positioned 1.5 mm lateral from Bregma and 0.2 mL of Rose Bengal solution (Sigma; 10g/L in saline, i.p.) was administered. After 5 min the brain was illuminated through the skull for 15 min. All animals were randomly assigned to a treatment group 5 days post-stroke, by an operator not undertaking behavioral, histological or immunohistochemical assessments. All assessments were conducted by observers blind to the treatment group.

### NMDA-induced excitotoxic damage

In isoflurane-anesthetized mice, excitotoxic lesions were induced by NMDA micro-injection (20 nmol/0.5 μL) into the right striatum (2.5 mm lateral, −4.0 mm ventral and −0.7 mm posterior to the Bregma) infused at a rate of 0.2 μL/min. The lesion volume was quantified 48 hours later.

### Peptide synthesis and drugs preparation

Mouse/rat ODN (H-Gln-Ala-Thr-Val-Gly-Asp-Val-Asn-Thr-Asp-Arg-Pro-Gly-Leu-Leu-Asp-Leu-Lys-OH) was synthesized as previously described (Leprince *et al*., 2001). Flumazenil, a selective antagonist of the benzodiazepine-site of the GABA_A_R and NMDA were purchased from Sigma-Aldrich and dissolved in sterile HEPES buffer supplemented with KCl (2.5 mmol/L) and NaCl (145 mmol/L) pH 7.4 and DMSO (dilution 1:4) for flumazenil. ODN or its vehicle was infused over 5 min into the lateral ventricle (3 μL; −0.1 mm posterior, 0.8 mm lateral, −2.5 mm ventral to the Bregma) or into the *cisterna magna* for the experiment conducted under the 2-photon microscope.

### *In vivo* dosing with hydrogel impregnated with ODN

A hyaluronan/heparan sulfate proteoglycan biopolymer hydrogel (HyStem^®^-C, BioTime, Inc.) was employed to locally deliver ODN, to the peri-infarct cortex as described previously (Clarkson *et al.,* 2011; Houlton *et al*., 2019). In brief, ODN was added to the HyStem/Gelin-S mix (component 1 of hydrogel, glycosyl/gelin 1:1), followed by addition of Extralink (component 2 of hydrogel) in a 4:1 ratio. In prior studies, we have reliably shown that hydrogels can be used to release small and large proteins for at least 3 weeks from the stroke cavity (Li *et al*., 2010; Overman *et al*., 2012). Five days after stroke, 7.5 μL of HyStem-C, impregnated with either ODN (1 μg or 5 μg) or saline-vehicle was injected directly into the stroke infarct cavity (30-gauge needle and a Hamilton syringe at coordinates 0 mm AP, 1.5 mm ML, and 0.75 mm DV).

### Behavioral tests

#### Middle cerebral artery occlusion

Behavioral tests were conducted by a manipulator blind to treatment groups. After a pre-test performed 3 or 4 days before MCAO surgery, 3 sensori-motor tasks were conducted once a week.

The *pole test* was performed as described by Matsuura *et al.,* 1997. Mice performed two different tasks: 1) mice were placed head upward on top of the vertical pole (55 cm). The time to turn completely head downward was measured (*return task*); 2) the descent of the pole (*descent task*) covered with Durapore tape (3M). The third test was the *beam crossing* test (Carter *et al*., 2001), measuring the time to cross a horizontal beam (diameter: 10 mm; length: 1 m). For animals unable to perform one of these 3 tasks, a time penalty of 60, 60 and 200 s was respectively attributed, corresponding to the worst performances (Mann and Chesselet, 2014).

#### Cortical photothrombosis

Recovery of forelimb motor function was determined by the cylinder and grid-walking tasks to assess their exploratory behavior and walking, respectively, as previously reported (Clarkson *et al*., 2010). Mice were tested approximately 7 days prior to stroke to establish a baseline performance level and then after 7, 14, 28 and 42 days post-stroke at approximately the same time each day. Observers blinded to the treatment group scored behaviors as previously described.

### Immunofluorescent labeling of GFAP and infarct volume

Brains were sectioned and every sixth section (30 μm thick) were collected and stored in cryopreservation solution. Slices were first incubated with a primary antibody (chicken anti-mouse GFAP, 1:3,000, AB5541, Millipore) for 48 hours at 4°C. Then, a secondary antibody (anti-chicken Alexa-488, 1:1,000, SA5-10071, ThermoFisher Scientific) followed by the nuclear counter stain Hoechst (1:1,000, Sigma-Aldrich) were used. Images were taken with an Olympus BX61 microscope. Changes in GFAP staining was investigated 2 and 6 weeks post-stroke, with measurements taken 0-200 μm and 800-1000 μm from the stroke border in layers 2/3 and 5. Using the software FIJI Image-J (NIH, USA) the integrated density value (IDV) was measured in all 4 regions of interest (ROIs).

Infarct volumes were determined 2 and 6 weeks post-stroke using cresyl violet staining and Image-J analysis. The analysis is based on obtaining measurements from every sixth section and infarct volume was quantified as follows: infarct volume (mm^3^) = areas (mm^2^) x (section thickness (mm) + section interval (mm)). Infarct volume were corrected for edema as described previously (Lin et al., 1993). All analyses were performed by an observer blind to the treatment groups.

### Whole-cell voltage-clamp electrophysiological recordings

Mice (2-3-month-old) were anesthetized with isoflurane 3-7 days post-stroke or sham surgery and brains were sliced and prepared for recordings of tonic currents. All recordings were made from peri-infarct pyramidal neurons within layer 2/3 of the primary motor cortex, as previously described (Clarkson *et al*., 2010, 2019). Neurons were voltage-clamped in whole-cell configuration using a MultiClamp-700B amplifier using microelectrodes (3–5 MΩ) filled with a cesium-methyl-sulfonate (CsMeSO_4_)-based internal pipette solution. The recording aCSF was supplemented with 5 μM GABA to replenish the extracellular GABA concentration reduced by the high-flow perfusion of the slices.

Tonic inhibitory currents (*I*_tonic_) were recorded as the reduction in baseline holding currents (*I*_hold_) after bath-applying a saturating amount (100 μM) of the GABA_A_R antagonist SR-95531 (gabazine), while voltage-clamping at +10 mV. ODN was added to the recording aCSF *via* perfusion and their effects on *I*_tonic_ were recorded as the post-drug shift in *I*_hold_. Drug perfusion continued until the shifting *I*_hold_ remained steady for 1-2 min.

### Statistics

Sample size was calculated using the JavaScript utilities available at www.stat.ubc.ca/~rollin/stats/ssize/index.html with parameters determined from prior work. For experiments presented in Figures 1A, 1D, 2B, 3D, the knowledge of the variability was too uncertain to reliably calculate a sample size (>80% power). For those experiments, we relied on reasonable assumptions to determine a priori, the number of animals used. Data are presented as mean±SEM or as box-and-whisker plot showing the median (box = first and third quartiles; whisker = range). Statistics were performed using Prism software (Graphpad). Normal distribution of the datasets was tested by a Kolmogorov-Smirnov test. Pairwise means comparisons were performed using *t*-test for normally distributed data, or Mann–Whitney test otherwise. For recovery studies, two-way ANOVA followed by post-hoc Tukey’s or Bonferroni’s test for multiple comparisons was performed.

**Figure 1.**
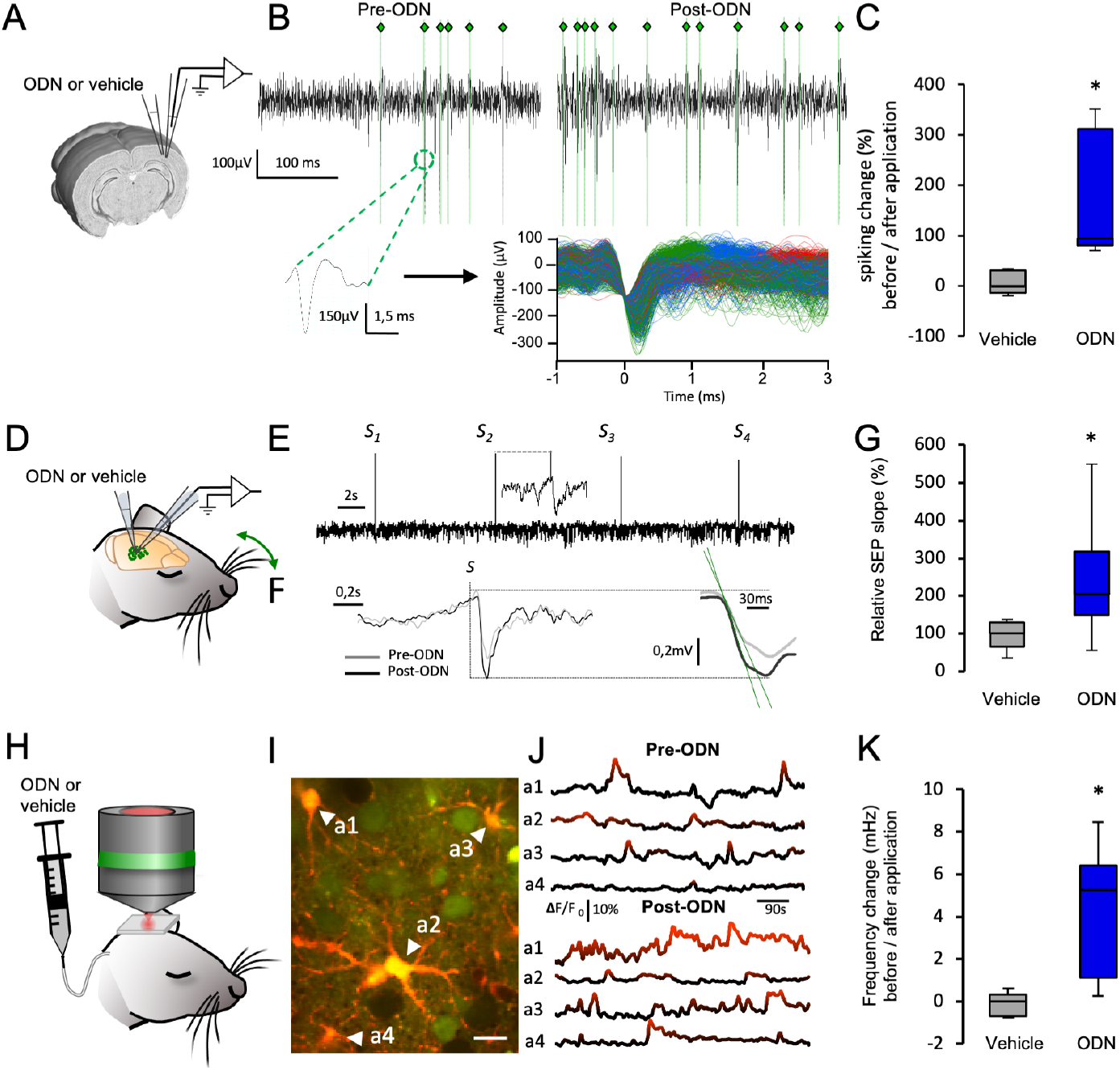
The gliopeptide ODN enhances neuronal and astrocytic activity in the cortex *in vivo*. **A**, Experimental arrangement showing pipette-positions for microinjection of ODN in the vicinity (~100 μm) of the recording pipette (layer 4). **B**, Up, representative extracellular unit recording trace comparing spontaneous neuronal spiking activity before and after ODN infusion (350 ms epoch). The post-treatment period recording started 6 min after the start of the infusion. The occurrence of spikes is depicted by vertical green lines underneath the trace. Down, representative examples of a detected spike (left) and a superimposed spike waveform showing 3 clusters of spikes (green, red and blue). **C**, The infusion of the vehicle solution did not change the number of spikes detected during the 12 min post-vehicle period (*P* > 0.05 vs. 12 min pre-vehicle period, n = 6 mice). ODN significantly increased spiking (*P* < 0.05 vs. pre-ODN period, n = 6 mice). **D**, Experimental set-up for sensory evoked potential (SEP) recorded in the whisker barrel cortex. The contralateral whisker pad was mechanically stimulated every 10 s evoking the typical negative shift of the LFP trace (**E)**. **F**, Representative SEP obtained after the average of 100 stimulations. **G**, The SEP slope of the control group remained unchanged after the infusion of vehicle (*P* > 0.05, pre- vs. post-treatment, n = 6 animals) whereas ODN treatment significantly increased the SEP slope (*P* < 0.05, pre- vs. post-treatment, n = 6 animals). **H,** Experimental set-up for 2-photon imaging of astrocyte activity. ODN or its vehicle was administered by the cisterna-magna route. **I**, Astrocytes (white arrowheads) were double-labeled with Ca^2+^-sensitive and Ca^2+^ insensitive dyes (OGB-1 and SR-101, respectively). Scale bar: 10 μm, depth: ~ −250 μm. **J**, Example of spontaneous somatic Ca^2+^ activity showing the occurrence of Ca^2+^ surges (red highlight). **K**, The infusion of the vehicle did not change the frequency of astrocytes Ca^2+^ transients (*P* > 0.05 vs. pre-vehicle period, n = 5 mice) whereas ODN significantly increased their frequency (*P* < 0.05 vs. pre-ODN period, n = 6 mice). Data are represented as box-and-whisker plot (box = first and third quartiles; whisker = range). Mean values were compared using paired, two-tailed t-test.

### Data availability

All the data that support the findings of this study are available from the corresponding author.

## Results

### Effect of ODN on cortical activity

Changes in neuronal excitation is critical for cell survival during the acute phase of stroke as well as for synaptic plasticity and recovery during the repair phase after stroke. To address the general hypothesis that extrinsic ODN can be used to safely manipulate neuronal excitability and influence post-ischemic repair, we first checked whether, as reported previously *in vitro*, ODN induced a measurable enhancement of cortical activity *in vivo*. In neuronal cell culture, the GABA_A_-R NAM effect of ODN was observed in the micro- to millimolar range (Guidotti et al., 1983; Ferrero et al., 1986; Alfonso et al., 2012). We therefore targeted this range to test ODN *in vivo*. In isoflurane-anesthetized mice, we recorded the effect of 1 μg of ODN or its vehicle on neuronal spiking of layer 4 of the somato-sensory cortex (Fig. 1A). Inspection of electrophysiological recordings did not reveal any aberrant oscillations or sharp events that would indicate epileptiform activity (Fig. 1B). ODN increased spontaneous neuronal firing (181.5±61.6% increase compared to the pre-treatment period; *P* < 0.05; n = 6 mice; vehicle: 4.7±11.4%; *P* > 0.05; n = 6) while vehicle administration did not result in any significant change in spiking activity (Fig. 1C). To further examine the potential of ODN to boost neuronal excitability, the effect of ODN was also tested during a somato-sensory stimulation. A single deflection of the whiskers pad triggered a somato-sensory evoked potential (SEP) in layer 4 of the contralateral barrel cortex (Fig. 1D-E). As a result of ODN treatment, SEP slopes increased by 147.8±71.8% showing that ODN increases neuronal excitability (*P* < 0.05 vs. pre-treatment period; n = 6 mice; vehicle group: 4.7±26.4%; *P* > 0.05 vs. pre-treatment period; n = 6; Fig. 1F-G). Furthermore, the positive effect of ODN on cortical activity was also revealed by the increase of astrocytic network activity seen by intravital calcium imaging using a two-photon microscope (Fig. 1H-J). Astrocytes displayed spontaneous calcium transients at 3.6±1.4 mHz during baseline as previously described (Chuquet *et al*., 2007). After ODN administration, transient frequency rose to 7.5±0.8 mHz (*P* < 0.05; n = 23 cells from 6 animals) whereas calcium activity was unaffected following vehicle administration (3.15±0.7 mHz vs. 3.08±0.8 mHz; *P* > 0.05; n = 16 cells from 5 animals; Fig. 1J-K). Overall these results show for the first time that the endozepine ODN acts as an excitability-enhancer of the cerebral cortex *in vivo*.

### Acute effect of ODN during cerebral ischemia

Within the first minutes to hours after stroke onset, depolarization and hyperexcitability are the primary culprits for the massive necrotic neuronal loss. Although ODN has a potent neuroprotective effect *in vitro* (Hamdi *et al*., 2011), enhancing cortical excitability in the acute phase of brain ischemia, when excitation is already lethal for neurons, may be inimical to cell survival. In order to examine whether the subtle changes of cortical activity elicited by ODN could influence ischemic neuronal death processes, we administered ODN (1 μg intracerebroventricular, i.c.v) during the acute phase of a focal brain ischemia elicited by intraluminal occlusion of the right middle cerebral artery (MCAO). The transient ischemia (60 min followed by reperfusion) produced a reproducible infarct 48 h later (Fig. 2A). Continuous laser Doppler flowmetry monitoring of the cerebral blood flow (CBF) confirmed that all animals underwent a similar ischemia and that ODN had no impact on residual CBF (*P* > 0.05 at any time point; Fig. 2C). The treatment with ODN resulted in a severe aggravation of the mean infarction volume (ODN group 140.35±14.61 mm^3^ vs. vehicle group 92.32±8,61mm^3^; *P* < 0.05; n = 8; Fig. 2A). Physiological parameters well known to have a determinant impact on neuronal survival in stroke (arterial pressure, temperature, food intake) were not affected by central administration of ODN (Fig. 2C-F). Flumazenil, a selective antagonist of the benzodiazepine-binding site of the GABA_A_R, fully reversed ODN-induced exacerbation (ODN+FLZ group 93.87±8.70 mm^3^ vs. vehicle group 92.32±8,61mm^3^; *P* > 0.05; n = 8; Fig. 2A) confirming the involvement of the GABAergic signaling. MCAO was also conducted on knockout mice for the DBI gene (DBI^−/−^) and consistent with the above observations, DBI^−/−^ animals appeared to be more resistant to MCAO than their wild-type (WT) counterparts (WT group 80.08±8.73 mm^3^ vs. DBI^−/−^ group 47.02±7.30 mm^3^; *P* < 0.05; n = 6; Fig. 2B). This result suggests that the endogenous production of the ODN precursor plays a role in the pathophysiology of stroke. In view of the above results, the most likely hypothesis for the ODN aggravated neuronal cell death, is due to a decrease in GABAergic inhibition, a well-documented pathway in neuroprotection and recovery (Green *et al*., 2000; Clarkson *et al*., 2010). This was further corroborated by two other observations. First, we examined the effect of ODN in the case of a pure excitotoxic neuronal cell death induced by a microinjection of NMDA. The co-administration of ODN with NMDA significantly increased the size of the excitotoxic lesion (3.5±0.4 mm^3^ vs. 6.4±0.9 mm^3^; *P* < 0.05; Fig. 3A-B). Second, another, albeit closely related pathophysiologic event, is spreading depressions (SD), *i.e.* waves of transient depolarization preceding cell damage in the compromised brain (Hartings *et al*., 2017). Local application of KCl on the healthy cortical tissue triggers such SD events, as described elsewhere (Chuquet *et al.,* 2007). We found that central administration of ODN 10 min before the induction of a train of 2-3 SDs, increased the number of events by 61.8% (*P* < 0.05 vs. vehicle group; n = 6; Fig. 3C-D). The amplitude and temporal features of SDs were unchanged by ODN (*P* > 0.05; n = 6; Fig. 3E). In a condition of hyperexcitability like the one observed during the acute phase of ischemia, the neuroprotective features of the gliopeptide ODN are surpassed by its pro-excitotoxic effect. Altogether, these observations are fully consistent with the view that the negative allosteric modulation of the GABA_A_R is not safe in the acute phase of stroke (Clarkson *et al*., 2010).

**Figure 2.**
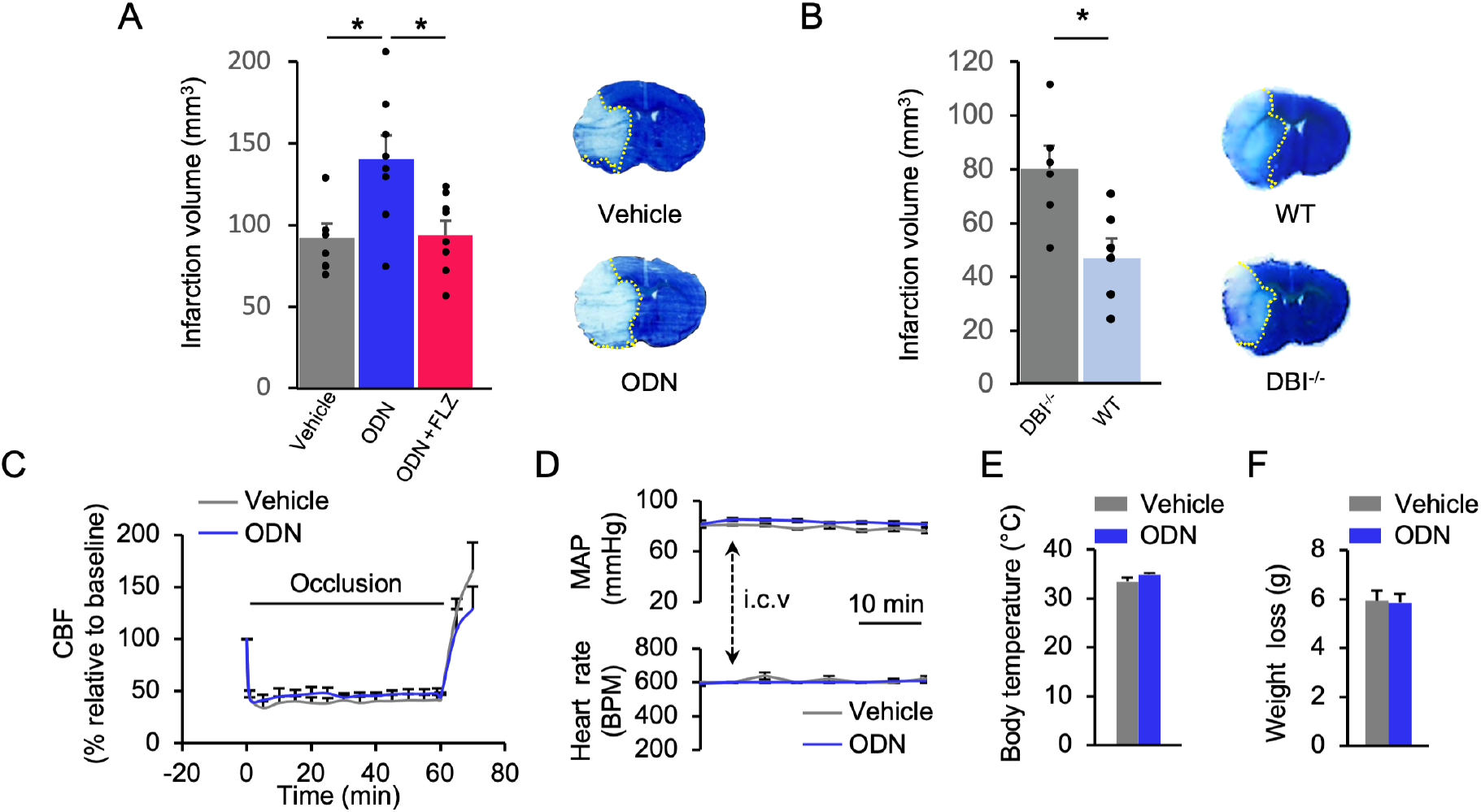
ODN exacerbates ischemic damage. **A**, Average infarction volume when treatments are provided during the acute phase of focal cerebral ischemia, by i.c.v injection. ODN significantly increased the volume of the lesion (*P* < 0.05 vs. vehicle group; n = 8 animals/group). This effect was reversed by the selective GABA_A_R benzodiazepine-site antagonist flumazenil (FLZ, *P* > 0.05 vs. vehicle group, n = 8 animals). Right, representative histological analysis showing the infarction, unstained by thionine. **B**, Knockout mice for DBI (DBI^−/−^, the peptide precursor of ODN) were less vulnerable than wild-type mice to brain focal ischemia (*P* < 0.05 vs. WT; n = 6 animals/group), representative lesions are shown on the right. **C-F**, The exacerbation of the damage could not be attributed to a difference in the severity of cerebral blood flow (CBF) decrease between groups during the procedure (**C**, *P* > 0.05 vs. vehicle, n = 8 animals/group). I.c.v administration of ODN produced no effect on systemic physiological parameters (**D**) such as mean arterial pressure (MAP, *P* > 0.05) or heart rate (beat *per* min., BPM, *P* > 0.05 vs. vehicle, vehicle n = 3; ODN n = 4), body temperature (**E**, *P* > 0.05 vs. vehicle, n = 8/group) or post-stroke weight loss (**F,***P* > 0.05 vs. vehicle, n = 8/group). Data are represented as mean±SEM. Mean values were compared using unpaired, two-tailed Mann-Whitney test (**A, B, E, F**) followed by Bonferroni’s post hoc testing for multiple comparison (**A**), or one-way ANOVA (**C, D**).

**Figure 3.**
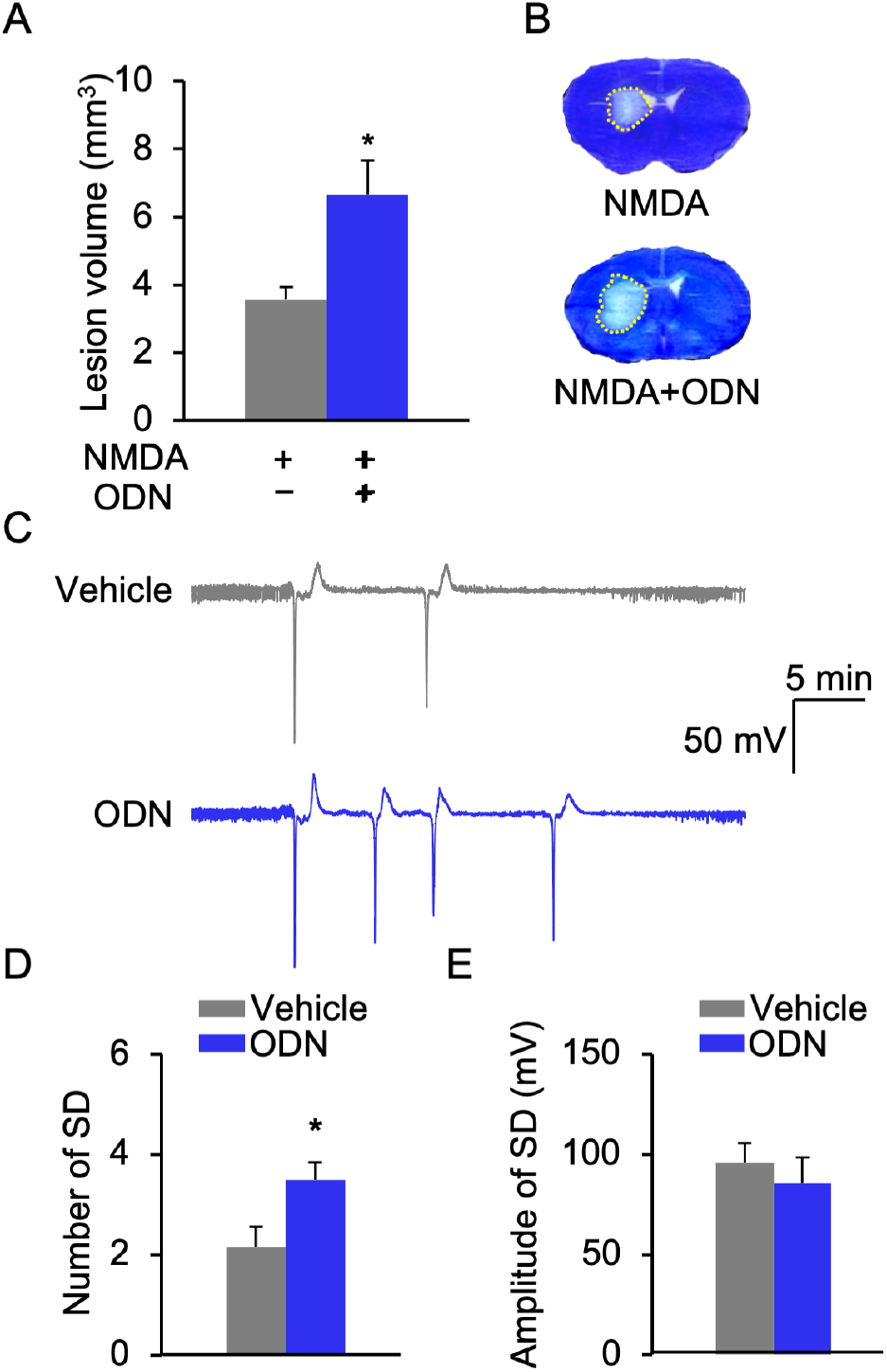
ODN enhances excitotoxicity and spreading depolarization waves. **A**, NMDA-induced striatal damages were exacerbated by co-administration of the gliopeptide ODN (*P* < 0.05; NMDA: n = 14 and NMDA+ODN: n = 13). **B**, Representative lesions are shown on the right. **C**, Intracortical infusion of KCl induced recurrent spreading depression (SD) waves: representative LFP traces showing the typical negative shift of SDs, whose number was increased in presence of ODN (*P* < 0.05, vehicle n = 6; ODN n = 6) while their amplitude remained unchanged (**D**, **E,***P* > 0.05). Data are represented as mean ± SEM. Mean values were compared using two-tailed Mann-Whitney test.

### Effect of ODN on functional recovery after MCAO stroke

In the weeks following stroke, the peri-infarct surviving tissue is in a state of heightened neuronal plasticity, which is intended to promote the recovery of lost functions. Some of the repair processes that are altered rely on an increase in synaptic transmission and therefore depend on an optimal excitation/inhibition balance. Contrary to the acute phase of stroke (hours) where excitation must be decreased to maintain neuronal survival, in the chronic phase (weeks), an excess of GABAergic inhibition prevents optimal recovery (Clarkson *et al*., 2010). Spurred by recent studies demonstrating that the exogenous benzodiazepine inverse agonist L655,708 is efficient in correcting this imbalance and improves the recovery of mice after stroke (Clarkson *et al*., 2010; Lake *et al.,* 2015), we made a similar hypothesis that chronic treatment with the endozepine ODN would promote enhanced neuronal plasticity. To avoid any interactions of ODN with its effects on cell death during the acute phase of stroke, we began the treatment with the gliopeptide (or its vehicle) 72 h after the onset of stroke (Fig. 4A-B), a sufficient delay for the lesion to be fully formed and no further expansion observed beyond this period. Preoperative performances in the “latency to turn” task, the “latency to descend” task and the “latency to cross” task did not differ between the four different groups. Altogether these three behavioral tasks provided a well-established method for measuring weekly evolution of sensorimotor coordination and balance following extended ischemic lesion (e.g. cortex and striatum) in rodents over a period of weeks following the initial insult (Matsuura *et al*., 1997; Carter *et al*., 2001). Unlesioned sham mice were not affected at any time-points, reporting that neither the surgery procedure, the ODN treatment, nor a learning effect interfered in the observations made from MCAO animals. Stroke induced a severe decrease in the ability to execute all three tasks (Fig. 4C-E), with only a small spontaneous gain of function observed over weeks 1 and 2 in the vehicle treated group. ODN treatment however, resulted in a marked improvement on all three task functions, with significance observed by weeks 3 and 4 post-stroke (Fig. 4C-E). Assessment on two of the three tasks revealed that the MCAO plus ODN treatment group was not significantly different when compared to unlesioned controls at day 28 post-stroke. ODN had no impact on stroke-induced weight loss, and mortality rate remained similar between ODN and vehicle-treated groups (Fig. 4F). At 28 days post-stroke, brains were collected for further histological analysis. The lesion characteristics *i.e* extent, cavities and ipsilateral atrophy, revealed no differences between MCAO-vehicle and MCAO-ODN groups, highlighting that ODN treatment was safe when administered from 3 days after the stroke onset (Fig. 4G-I).

**Figure 4.**
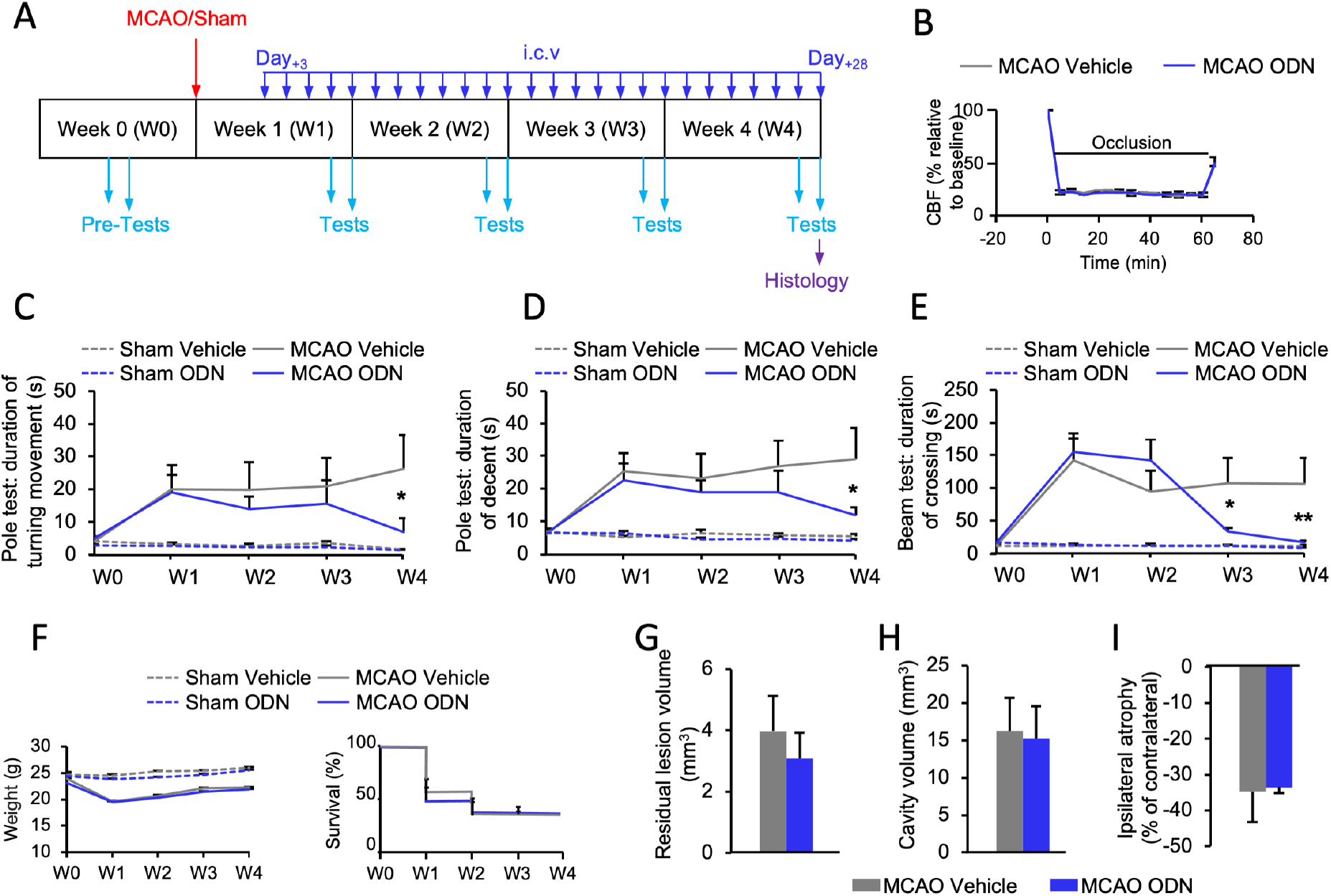
Delayed and chronic treatment with ODN promotes functional recovery after stroke in young adult male mice. **A**, Time-line diagram of the protocol. Treatment started 3 days after the middle cerebral artery occlusion (MCAO) procedure (MCAO ODN, n = 7; MCAO vehicle n = 6) or the surgery procedure without the MCAO (sham ODN, n = 6 and sham vehicle, n = 6). ODN or its vehicle were i.c.v injected everyday during the next 25 days (dark blue arrows). Behavioral tests were carried out at the end of each week (light blue arrows). MCAO was similar in ODN treated group and vehicle treated group (**B**). Mice treated with ODN progressively improved their performances in the execution of (**C**) the turning movement at the top of the vertical pole (*P* < 0.05 at week 4 relative to vehicle treated animals), (**D**) the descent of the vertical pole (*P* < 0.05 at week 4 relative to vehicle treated animals) and (**E**) the crossing of the horizontal beam (*P* < 0.05 at week 3 and *P* < 0.01 at week 4 relative to vehicle treated animals). Sham animals were not sensitive to ODN at any time point, and neither the surgical procedure nor the time affected their performance. Weight loss (**F,** left, *P* > 0.05 vs. MCAO+vehicle or Sham+vehicle; MCAO+vehicle n = 6; MCAO+ODN, n = 7; Sham+vehicle n = 6; Sham+ODN, n = 6) and mortality (**F** right, Logrank Mantel-cox test *P* > 0.05 vs. MCAO+vehicle, n = 6; MCAO+ODN, n = 7) were not different between groups at any time point. At the end of the procedure, histological analysis of mouse brain showed no difference between groups for the size of the residual lesion (**G,***P* > 0.05 vs. MCAO+vehicle; MCAO+vehicle n = 6; MCAO+ODN, n = 7), the total volume of cavities (**H**) and the atrophy of the ipsilateral hemisphere (**I**, *P* > 0.05 vs. MCAO+vehicle; MCAO+vehicle n = 6; MCAO+ODN, n = 7). Data are represented as mean ± SEM. Data were compared using Mann-Whitney test (**G**, **H**, and **I**) or nonparametric 2-way ANOVA followed by Bonferroni’s post hoc testing (**B**, **F** left, **C**, **D**, **E**).

### Effect of ODN on functional recovery after photothrombotic stroke

Although the intra-arterial filament occlusion is the gold-standard model for pre-clinical investigation, a way to strengthen the preclinical evidence of a drug is to challenge it in multiple experimental paradigms. Indeed, the intra-arterial filament occlusion model results in a large corticostriatal infarction, mimicking malignant infarction, while human strokes are mostly small in size. The photothrombotic model produces a small cortical infarction with well-delimited boundaries useful for modeling post-stroke impairments (Corbett et al., 2017). Furthermore, this model is well suited for hydrogel-delivery of peptides and small proteins diffusing over several weeks from the stroke cavity into peri-infarct tissue (Overman et al., 2012; Clarkson et al., 2015; Cook et al., 2017). To further differentiate this second model and comply with stroke basic research recommendation (Bernhardt et al., 2017) we used aged female mice (20±2 months) representing a population of stroke sufferers with more severe damage and poorer recovery after the event compared to aged men (Koellhoffer and McCullough 2013). Hydrogel impregnated with ODN was one-time injected into the infarction cavity 5 days after the onset of stroke (Fig. 5A). In this protocol of progressive diffusion from a single depot of gel (7.5 μL), 2 doses of ODN were tested (1 and 5 μg). Photothrombosis produced a well-circumscribed cortical lesion of 3.68±0.43 mm^3^ (n = 5 mice) as assessed 2 weeks post-stroke, that resulted in some spontaneous resolution over time with the infarct volume being 2.9±0.66 mm^3^ (n = 10 mice) as assessed 6 weeks post-stroke at the completion of the behavioural experiments. Treatment with ODN resulted in a dose-dependent decrease in infarct volume (F_(2,40)_ = 6.742; *P* < 0.003: ODN (1 μg), 3.44±0.54 mm^3^, and 1.86±0.50 mm^3^ for weeks 2 and 6, respectively; ODN (5 μg), 2.73 ± 0.84 mm^3^, and 1.51±0.39 mm^3^ *P* < 0.05, for weeks 2 and 6, respectively; Fig. 5B-C).

**Figure 5.**
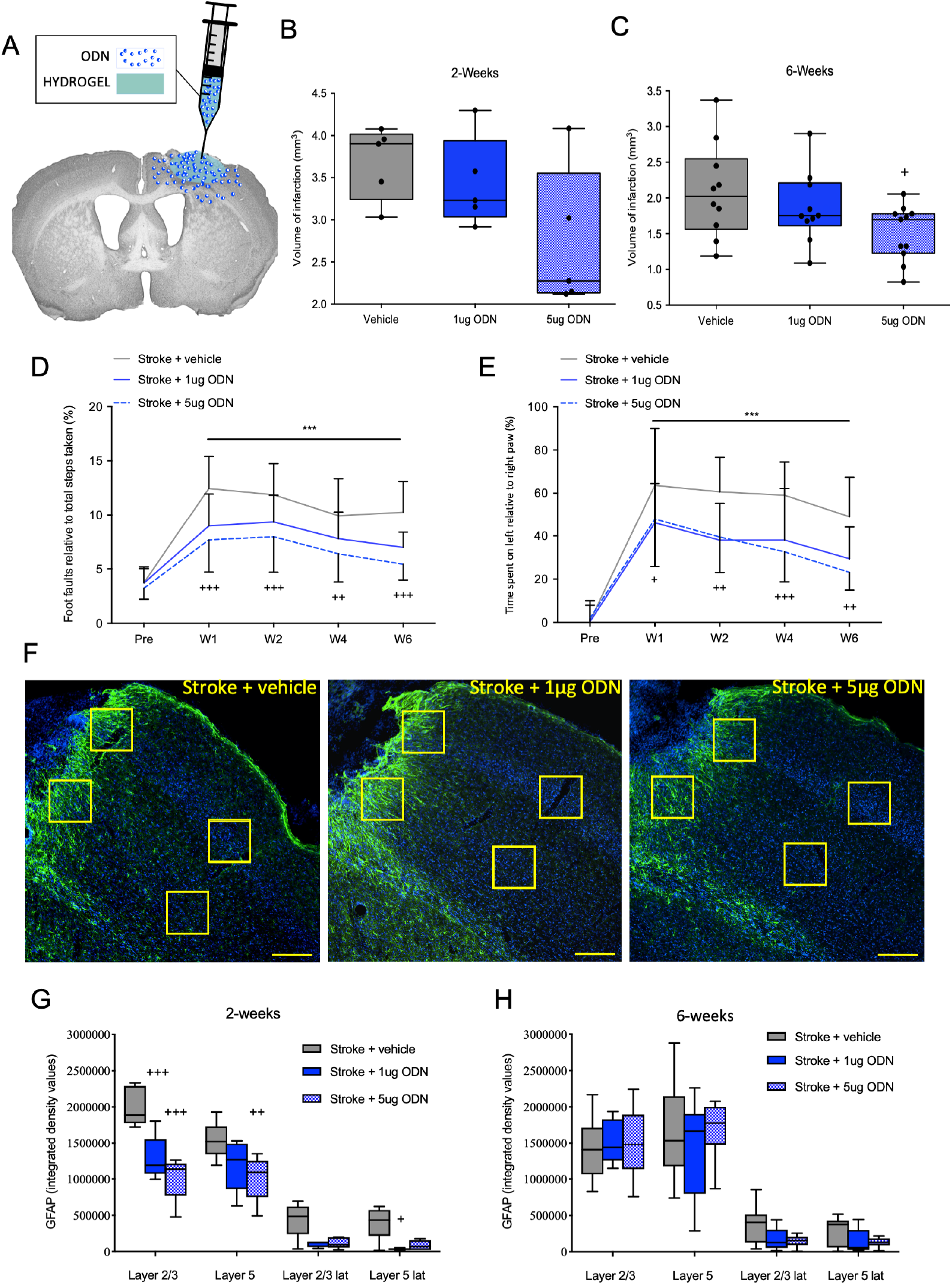
Delayed, chronic treatment with ODN-impregnated hydrogel promotes functional recovery after stroke in aged female mice. **A**. Schematic illustration of the injectable ODN-impregnated hydrogel. Infarct volume was assessed at 2 (**B**) and 6 (**C**) weeks post-stroke using cresyl violet staining (n = 5 and 10 respectively). Assessment of 1 μg ODN-treated animals showed no differences in infarct volume compared to vehicle treated controls. However, 5 μg ODN-treated animals exhibited a progressive decrease in stroke volumes with infarct volume being significantly different to vehicle-treated stroke controls 6 weeks post-stroke (^+^ *P* < 0.05; n = 10 animals/group). Motor behavioral function was assessed using both the gridwalking (**D**) and cylinder tasks (**E**). On both tasks, ODN (both 1 and 5 μg) resulted in a decrease in the number of footfaults on the gridwalking task and an improvement in forelimb asymmetry in the cylinder task. As ODN is a gliopeptide with astrocyte protective features, we examine whether ODN treatment affected the extent of peri-infarct glial scaring as assessed by GFAP staining (**F-H**). Representative GFAP strained sections are shown for stroke + vehicle (**F, left**), stroke + 1 μg ODN (**F, middle**) and stroke + 5 μg ODN (**F, right**) treated animals, 2 weeks post-stroke (scale bar is 200 μm). GFAP staining intensity was assessed from layers 2/3 and 5, 0-200 μm and 800-1000 μm (each box represents 200 x 200 μm) from the stroke border at 2 (**G**) and 6 weeks (**H**) post-stroke. Treatment with ODN resulted in a dose-dependent decrease in reactive astrogliosis as assessed by GFAP-labeling 2 weeks post-stroke (**G**). No differences in GFAP-labelling were observed at 6 weeks post-stroke between treatment groups (**H**). ^+++^ *P* < 0.001, compared to pre-stroke baseline behavioural controls; ^+^ *P* < 0.05, ^++^ *P* < 0.01, ^+++^ *P* < 0.001, compared to stroke + vehicle controls.

We next tested the mice behaviorally on both the gridwalking (forelimb function) and cylinder (forelimb asymmetry) tasks. Behavioral assessments revealed an increase in the number of footfaults on the gridwalking test (n = 10 per group; Fig. 5D) and an increase in spontaneous forelimb asymmetry in the cylinder task (n = 10 per group; Fig. 5E) from one-week post-stroke. ODN treatment resulted in a dose-dependent decrease in the number of footfaults on the gridwalking task (time effect: F_(4,150)_ = 33.56, *P* < 0.0001; treatment effect: F_(2,150)_ = 28.22, *P* < 0.0001) and an improvement in forelimb asymmetry in the cylinder task (time effect: F_(4,150)_ = 48.54, *P* < 0.0001; treatment effect: F_(2,150)_ = 18.73, *P* < 0.0001).

### Effects of ODN on modulation of the glial scar

As ODN is a gliopeptide with astrocyte protective feature (Hamdi *et al*., 2011), we next investigated whether exogenous administration of ODN has an effect on the glial scar. The glial scar has many roles after a brain ischemia has occurred, both as a regulator of inflammation, but also as a regulator of axonal sprouting and brain excitability, which are critical processes for cortical remapping (Clarkson *et al*., 2010; Sofroniew, 2015; Anderson *et al.*, 2016; Brown et al., 2019; Lie *et al.,* 2019). We observed a clear up-regulation in glial fibrillary acidic protein (GFAP) expression in the peri-infarct region at both 2 and 6 weeks post-stroke, with expression levels decreasing further away from the stroke border we examined (Fig. 5F-H). Chronic treatment with ODN resulted in a dose-dependent decrease in the expression of GFAP in the peri-infarct in both layers 2/3 (both 1 μg, *P* < 0.001; and 5 μg, *P* < 0.001), and 5 (only for 5 μg, *P* < 0.001) as assessed 2 weeks post-stroke (Fig. 5G). Assessment of GFAP expression in peri-infarct regions 6 weeks post-stroke revealed no differences in expression between vehicle and ODN treated animals in either layers 2/3 or 5 (Fig. 5H).

### Effects of ODN on tonic inhibitory currents

Given the profile of motor functional recovery we next sought to investigate the changes in tonic inhibitory currents following treatment with ODN. This is also driven by the fact that we have previously reported that tonic inhibitory currents are elevated from 24 h after stroke and remain elevated for extended period of time due in part to the formation of the glial scar (Clarkson *et al*., 2010, 2019; Lie *et al*., 2019). Tonic inhibitory currents were assessed in the peri-infarct cortex of mice after a photothrombotic stroke to the forelimb motor cortex. Layer 2/3 pyramidal neuron whole-cell voltage-clamp recordings were obtained from brain slices generated *ex vivo* 3-7 days after stroke (Fig. 6A-B). Recordings obtained 3-7 days after stroke showed an increase in GABA_A_R-mediated tonic inhibition (Control: 53.13±20.04 pA vs. Stroke: 130.90±39.01 pA; *P* < 0.001; Fig. 6B). As ODN is released exclusively by astrocytes, extrasynaptic GABA_A_ receptors are a likely candidate to mediate its effect on neuronal excitability (Tonon *et al*., 2020). Therefore, we next examined if bath application of ODN could correct peri-infarct tonic inhibition. Bath application of ODN to slices generated 3-7 days post-stroke resulted in a significant decrease in GABA_A_R-mediated tonic inhibition (Stroke + ODN: 91.25±13.49 pA; *P* < 0.05; Fig. 6B) compared to stroke controls. It should be noted however, that this decrease in tonic inhibitory currents was only a partial reversal with tonic inhibitory currents still significantly elevated compared to sham controls (*P* < 0.05).

**Figure 6.**
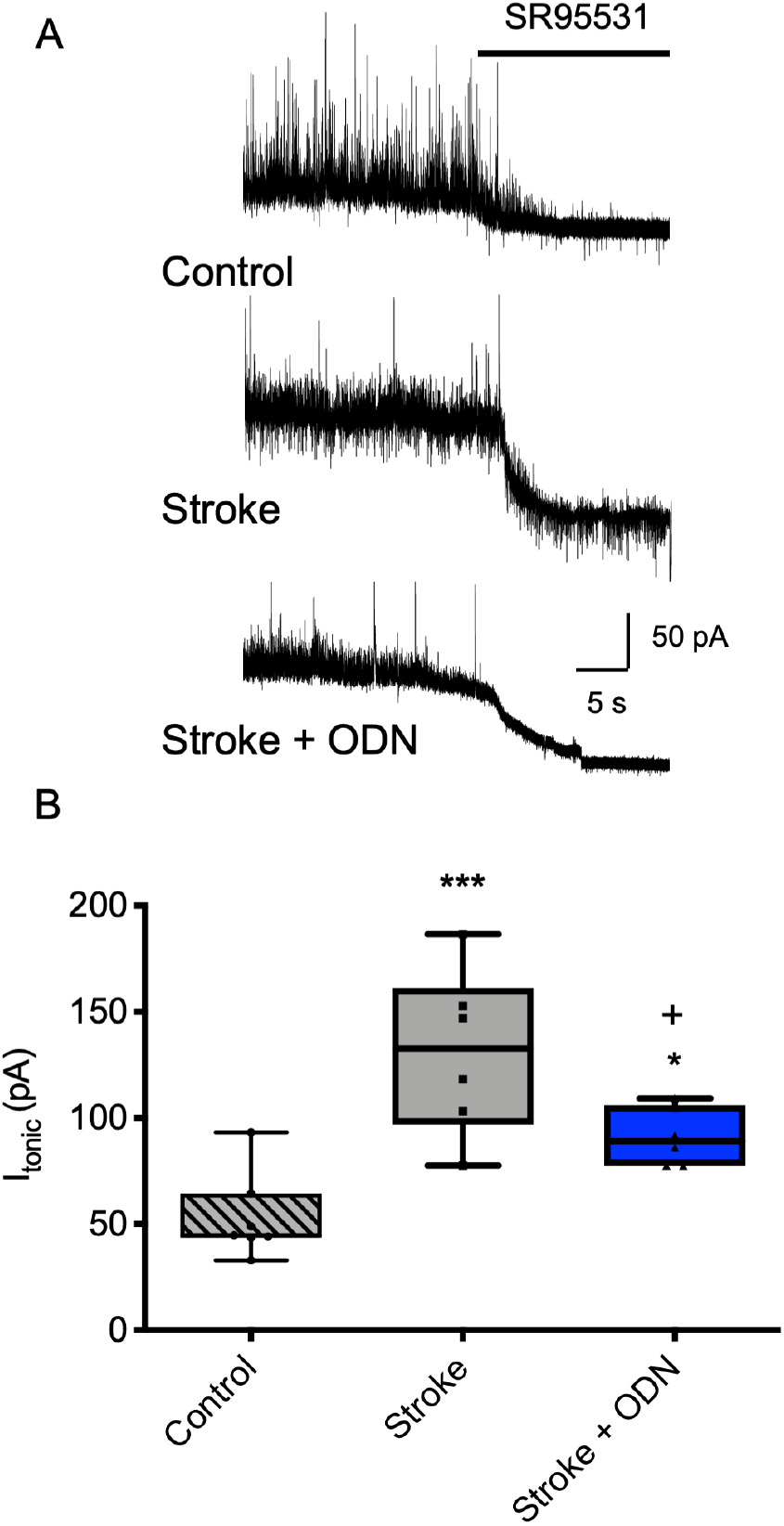
ODN dampens the stroke-induced elevation in tonic GABA_A_R currents in layer 2/3 pyramidal neurons. Whole-cell patch-clamp recordings were made from post-stroke brain slices, within 500 μm of infarct, from layer-2/3 pyramidal neurons. Representative traces showing the tonic inhibitory currents in control (n = 7), stroke (n = 6) and stroke + ODN (n = 6) treated animals (**A**). Tonic currents were revealed by the shift in holding currents after blocking all GABA_A_Rs with gabazine (SR95531; 100 μM). Box plot (whiskers: minimum and maximum; lines: median) showing an increase in tonic inhibitory currents in peri-infarct cortical neurons that is partially dampened following exposure to ODN (**B**). Horizontal bar indicates the application of the GABA_A_R antagonist, SR95531. Cells were voltage-clamped at +10mV. ^*^ *P* < 0.05, ^***^ *P* < 0.001, compared to tonic currents from control animal; ^+^ *P* < 0.05, compared to tonic currents from stroke animal.

## Discussion

Studies over recent years have shown that the peri-infarct cortex, which is the tissue adjacent to the stroke, is in a state of heightened plasticity during the sub-acute period (Murphy and Corbett, 2009). The process of plasticity is highly influenced by GABA_A_R signaling and tonic inhibition, which maintains and shapes the level of neuronal excitability (Raimondo *et al*., 2017). Alteration in GABAergic signaling has been previously reported to be triggered by stroke in both animals and humans (Carmichael 2012; Johnstone *et al*., 2018). Accordingly, dampening tonic inhibitory currents using a NAM targeting α_5-_containing GABA_A_Rs from 3-7 days post-stroke, but not acutely within hours, increases motor functional recovery in animal stroke models (Clarkson *et al*., 2010; Lake *et al.,* 2015). Consistent with these observations, the present findings demonstrate that exogenous administration of ODN enhances cortical excitability in a magnitude that makes it toxic when administered during the acute stage of stroke (within hours of stroke onset), however, makes it instrumental to safely boost functional recovery when administered during the sub-acute (3-7 days) recovery stages of stroke.

The endozepine, ODN, has been repeatedly categorized as a GABA_A_R NAM (Bormann *et al.,* 1985; Costa and Guidotti, 1991; Alfonso *et al*., 2012; Dumitru *et al*., 2017). Yet, all these experiments have been conducted *in vitro*. Here, we report for the first time, *in vivo,* that ODN increases cortical neuronal activity and excitability. Interestingly, we also report that astrocytes respond to the treatment. The question of whether this activation is a consequence of neuronal activity or a direct effect on astrocytes, as we previously report in cell culture, remains unclear (Gach *et al*., 2015). However, it is possible that ODN works on both systems as we report herein that administration of ODN partially dampens both the stroke-induced elevation in tonic inhibitory currents as well as the level of reactive astrogliosis in the peri-infarct cortex.

Capitalizing upon our confirmation of ODN as an enhancer of cortical excitability we assessed the effect of a gain or loss of ODN during the acute phase of brain ischemia/reperfusion. In line with our predictions, when ODN was administrated during brain ischemia, cerebral damage was dramatically increased. In addition, the finding that DBI^−/−^ mice were less vulnerable to transient focal ischemia than wild-type mice, suggests that even the endogenous production of DBI (and therefore ODN) during stroke, is deleterious.

In the acute stage of stroke, minor changes in various physiological and metabolic parameters can have major consequences on tissue viability. We first verified that i.c.v administration of ODN did not change body temperature, food intake or cardiovascular activity. Second, it should be noted that ODN is not pro-apoptotic or pro-inflammatory (Hamdi et al., 2011; Kaddour et al., 2013). Third, although the quality of the brain microcirculation is critical in the penumbra, we did not find any evidence of a vasomotor activity of ODN, such as vasoconstrictions of brain arteries (our personal observations). Although GABA_A_R are expressed in many cell types, only those containing the gamma2 subunit can bind BZ and potentially ODN. Indeed, this subunit is very frequent in neurons but weakly or not expressed in oligodendrocytes, microglia, astrocytes and endothelial cells of adult brain mice (Cahoy et al., 2008). We are left, then, with a primarily effect of ODN on inhibitory transmission and the most parsimonious explanation for why ODN exacerbated the cellular damage following stroke is due to an increase in neuronal excitation. This is in line with the effect of the GABA_A_R NAM L655,708 when used too early after a cortical ischemia (Clarkson et al. 2010). In support of this view, we report here that ODN exacerbates NMDA induced excitotoxic damage. We also show that ODN increases the number of spreading depolarizations waves (SDW). The trigger of SDW is actually settled over the level of neuronal excitation and their propagation out from the site of infarction is known to be a leading cause of ischemic infarct growth (Chuquet *et al*., 2007; von Bornstädt *et al*., 2015). Therefore, for neurons at immediate threat of death during the acute phase (< 3 days), ODN may amplify the depolarization and lead to an irreversible calcium overload. Our results indicate that there is a need to develop specific endozepine blockers to prevent their action on GABA_A_Rs during the acute phase of cerebral ischemia.

To date, no NAMs that target the benzodiazepine-binding site to reduce GABAergic activity and that lack any pro-convulsant or anxiogenic effects are available for clinical use. However, clinical trials using α5-containing GABA_A_R NAMs for enhancing stroke recovery are being explored in phase II trials (ClinicalTrials.gov ID: Hoffman La-Roche - NCT02928393; Servier RESTORE BRAIN Study - NCT02877615). It is increasingly recognized that GABAergic NAMs used at a safe dosage and that target specific subunit compositions, will be therapeutically useful, for example to improve cognitive functions in brain conditions where diminished excitability has to be boosted (Bolognani *et al*., 2015; Soh and Lynch, 2015). There is also growing evidence that impairment in GABA transporter (GAT-3/GAT-4) function contributes to the peri-infarct zone changes in neuronal inhibition and post-stroke functional recovery (Clarkson *et al*., 2010, Carmichael, 2012). As a consequence, the excessive GABAergic tone inhibits sensorimotor recovery and is likely due in part to synaptic plasticity and long-term potentiation (LTP), being sub-optimal (Atack *et al*., 2006). Two independent studies have demonstrated that the use of a benzodiazepine-site specific NAM improves post-stroke motor recovery in adult males (Clarkson *et al*., 2010; Lake *et al.,* 2015). In addition, a recent study showed that the treatment with the GAT3 substrate, L-isoserine, administered directly into the stroke cavity, can increase GAT3 expression and improve functional recovery after focal ischemic stroke (Lie *et al.,* 2019). Following this idea and protocol, we found that daily administration of ODN started 3 days after ischemia or hydrogel delivery from 5 days (*i.e*. when the risk of ODN induced cell death had ceased) remarkably improved motor coordination. Of note, the precursor of ODN, DBI, is one of the most transcripted genes in astrocytes (Zhang *et al*., 2014). The fact that GABA inhibits the release of ODN by astrocytes (Patte *et al*., 1999), suggests that in the neighbourhood of the stroke lesion, the excess in ambient GABA is likely preventing sufficient endozepine production.

This functional recovery cannot be due to a difference in stroke severity between groups because *i)* the average decrease of blood flow was identical over the 60 minutes of ischemia, *ii)* the mean brain histological sequelae were similar at day 28 and *iii)* mortality rate had the same kinetics in both groups. Moreover, ODN treated animals did not display any obvious behavior indicative of anxiety, aggression or hyperactivity suggesting that both the timing and dose of ODN administered led to a safe and efficient treatment to promote functional recovery.

To date, all translational studies have failed to treat stroke in part because preclinical studies routinely failed to utilize clinically relevant animal models. Several recommendations have been made, among which were the use of aged animals of both sex since aging remains the major non-modifiable risk factor for stroke (Jolkkonen and Kwakkel 2016; Corbett et al., 2017). Aged females represent the greatest percentage of stroke sufferers, with strokes in this population being more severe and recovery being worse than males and younger females (Koellhoffer and McCullogh 2013). Therefore, to challenge ODN efficacy and increase the translational value of our results, we studied the effect of ODN in a radically different experimental paradigm. Using the photothrombotic model of ischemia in aged female mice, we obtained a small infarction circumscribed to the cortex. We took advantage of the topology of the lesion to also test whether a single depot of ODN into the infarction cavity on day 5 post-stroke would improve functional recovery. Although the number of new peptides entering clinical trials continues to grow, their therapeutic potential in neurology has not yet been realized because they rarely get through the blood-brain-barrier and their half-life *in vivo* limits the time window to exert their action (Penchala *et al*., 2015). Hydrogel-impregnated with peptides or small proteins has been extensively used as a system of chronical delivery (for review, Fernandez-Serra 2020). Specifically, we and others have shown that biomolecules embedded into hydrogels have a sustained, consistent release into surrounding tissue lasting around 3-4 weeks (Tae et al., 2006; Li et al., 2010; Overman et al., 2012; Clarkson et al., 2015; Cook et al., 2017). For instance, cyclosporin, a 11-amino acids long peptide of 1,2 kDa (i.e. similar to ODN, 18-residue long for 1,9kDa) is slowly released by the hydrogel in a 2 mm cortical zone around the gel for at least 24 days (Tuladhar et al., 2015). Therefore, the hydrogel provides a protective micro-environment for ODN and allows its controlled delivery around the lesion, where excitability need to be enhanced. In this second stroke model, ODN was able to improve sensory-motor task performance as soon as week 1. These studies highlight a novel example for hydrogel delivery systems offering a support for effective peptide biodisponibility that bypasses the blood-brain-barrier to achieve consistent recovery post-stroke. Although 1μg was effective in the MCAO protocol, only the dose of 5μg was effective in the photothrombotic protocol. This is not surprising considering that in the MCAO protocol, a daily dose of 1 μg was administrated, while in the photothrombotic protocol, ODN was delivered to the peri-infarct cortex from a single depot of hydrogel. Furthermore, others have also observed that, when hydrogel is used to vehiculate a bioactive molecule in the brain, such as neurotrophins, a higher quantity of the later is necessary (Ravina et al. 2018).

Overall, our data are strikingly reminiscent of the findings of Clarkson *et al.,* 2010, and Lake *et al.,* 2015. Using a small naturally occurring peptide that is expressed and released by astrocytes and known to act as a GABA_A_R NAM, we found the same versatile effect in stroke as with the synthetic GABA_A_R NAM, L655-708: deleterious in the acute phase but promotor of recovery in the chronic phase. L655-708 is an imidazobenzodiazepine selective for the extrasynaptic GABA_A_R that contains the benzodiazepine sensitive subunit α_5_ and, as an inverse agonist, reduces the tonic inhibition. Interestingly, L655-708 has been shown to facilitate LTP by shifting toward less inhibitory activity (Atack *et al*., 2006). Similarly, we found that ODN was able to dampen the ambient tonic inhibition surrounding the stroke lesion. This disinhibition may be permissive for plastic rearrangements in the peri-infarct cortex to occur (Ziemann *et al.* 2001; Cicinelli *et al.,* 2003; Clarkson and Carmichael, 2009). Therefore, one hypothesis requiring future attention is that ODN selectively binds to α_5_ containing GABA_A_R. It is also possible that these changes in tonic inhibitory currents could be occurring through changes in presynaptic GABA signalling or astrocytic function, which is another two suggested pathways that need to be explore in furture experiments. In parallel, another possible mechanism that needs to be examined is neurogenesis. The group of Monyer have recently shown that ODN reduces the GABA-mediated current of neural stem/progenitor cell of the SVZ, inducing their proliferation and migration (Alfonso et al., 2012; Dumitru et al., 2017) supporting a role of endozepines in neurogenesis. Additional work should be undertaken to test the hypothesis that post-stroke neurogenesis can be controlled by ODN to optimize functional recovery.

The modification of network connections and synaptic strength allowing functional recovery requires that homeostatic mechanisms, such as the maintenance of excitation/inhibition balance at the network, single cell and synapse level, be in their operating range (Buzsáki, 2007). Our results suggest that endozepines play a significant role to tuning this balance and reinforce the idea that an appropriate correction of GABAergic tone at the right time can facilitate functional recovery after stroke.

## Acknowledgement

We thank Dr Arnaud Arabo who provided help and expertise that greatly assisted the research.

